# ERGA-BGE Reference Genome of the Northern chamois (*Rupicapra rupicapra*): Europe’s most abundant mountain ungulate

**DOI:** 10.1101/2024.11.20.624344

**Authors:** Elena Buzan, Aja Bončina, Boštjan Pokorny, Nuria Escudero, Rosa Fernández, Astrid Böhne, Rita Monteiro, Marta Gut, Francisco Câmara Ferreira, Fernando Cruz, Jèssica Gómez-Garrido, Tyler S. Alioto, Leanne Haggerty, Fergal Martin, Diego De Panis

## Abstract

The reference genome of *Rupicapra rupicapra* (subsp. *rupicapra*) provides insights into the genetic makeup that enabled this iconic mountain ungulate to adapt to its harsh environment, including its ability to survive in extreme weather and high altitudes—factors that are increasingly important in the face of climate change. A total of 29 contiguous chromosomal pseudomolecules were assembled from the genome sequence. This chromosome-level assembly encompasses 2.62 Gb, composed of 124 contigs and 76 scaffolds, with contig and scaffold N50 values of 77 Mb and 101 Mb, respectively.

## Introduction

The chamois, genus *Rupicapra*, belongs to the order Artiodactyla, family Bovidae, subfamily Caprinae, tribe Rupicaprini (Hassanin et al., 2009). Currently, the most common taxonomic classification considers the existence of two species within the genus *Rupicapra*, based on a combination of morphometric, genetic, and behavioural characteristics: the northern chamois (*Rupicapra rupicapra*) and the southern chamois (*R. pyrenaica*); they are further subdivided into seven (*rupicapra, balcanica, cartusiana, carpatica, caucasica, asiatica, tatrica*) and three (*pyrenaica, ornata, parva*) subspecies, respectively (Corlatti et al., 2011). The chamois is the most abundant mountain-dwelling ungulate in Europe and the Near East, and has also been introduced to New Zealand. Chamois of all subspecies are nearly monomorphic, except for a seasonal sexual dimorphism in body mass before the rut in late autumn, when males are notably heavier than females (Rughetti & Festa-Bianchet, 2010).

The most abundant subspecies, the Alpine chamois (*R. r. rupicapra*), is distributed throughout the Alps and is assessed as of least concern by the IUCN Red List (www.iucnredlist.org/species/39255/195863093).

The other subspecies are only present in smaller areas where they face several threats, including poaching, overhunting, human disturbance, competition with livestock, habitat loss and degradation, infectious diseases, hybridization with introduced individuals of other subspecies, and climate change (Corlatti et al., 2011). Although the conservation status of the genus *Rupicapra* is generally favourable (both the northern and the southern species are abundant), five subspecies are threatened (*R. r. balcanica* and *R. r. tatrica* are critically endangered), three are decreasing in population size or range, and even some populations of the most abundant subspecies (including *R. r. rupicapra*) have started to decline (Anderwald et al., 2020). The importance of environmental heterogeneity in shaping chamois’ behavioural and life history traits, especially in response to ongoing climatic change is increasingly evident. This will be one of the biggest challenges for the future of chamois research and conservation.

The range of the northern (Alpine) chamois in Slovenia, from which individuals were sampled for producing the reference genome, encompasses three different regions, each with a unique topography, habitat connectivity, interspecies interactions, and abundance of chamois: the Alps, Dinaric, and the Pohorje mountains. The chamois habitat is extensive and more or less continuous in the Alps, but suboptimal and fragmented in the remaining regions, including Pohorje Mts., where we collected samples for the reference genome. Importantly, in the 19th century, the chamois was reduced to dangerously low numbers in the Alps by hunting and was almost extirpated in the Dinaric Mts. (Buzan et al., 2013). In the late 1970s, chamois populations in the Eastern Alps, including Slovenia, were badly decimated by catastrophic outbreaks of sarcoptic mange epidemics. However, the Pohorje population was not affected by the disease outbreak (Apollonio et al., 2010).

Climate change has a significant impact on mountain biodiversity, including species and populations as well as their underlying gene pools. Animal populations inhabiting mountain habitats are especially vulnerable. Although suitable climates still exist at higher altitudes, they are expected to be restricted to ever smaller areas in the future. At the species scale, the Northern chamois occupies a large diversity of habitat types, from low-elevation forested areas to high-altitude alpine meadows, or where the terrain is steep and rocky. For this reason, the chamois represents an ideal model species to study potential evolutionary responses to environmental change, in particular mechanisms of local adaptation to climatic conditions.

With the availability of a reference genome, we can understand the Northern Chamois’s genomic background. This understanding allows us to detect and comprehend hybridization events and risks within selected populations, evaluate populations’ resilience in the face of environmental change, and understand the factors that influence their potential to persist and adapt. Addressing these key aspects is essential for effective conservation, adjusting management strategies, and ultimately ensuring the long-term survival of the chamois.

The generation of this reference resource was coordinated by the European Reference Genome Atlas (ERGA) initiative’s Biodiversity Genomics Europe (BGE) project, supporting ERGA’s aims of promoting transnational cooperation to promote advances in the application of genomics technologies to protect and restore biodiversity (Mazzoni et al., 2023).

## Materials & Methods

ERGA’s sequencing strategy includes Oxford Nanopore Technology (ONT) and/or Pacific Biosciences (PacBio) for long-read sequencing, along with Hi-C sequencing for chromosomal architecture, Illumina Paired-End (PE) for polishing (i.e. recommended for ONT-only assemblies), and RNA sequencing for transcriptomic profiling, to facilitate genome assembly and annotation.

### Sample and Sampling Information

On November 8, 2022, an adult, 3-year-old male of *Rupicapra rupicapra* was sampled by Elena Buzan and Boštjan Pokorny. The species identification through COI barcoding was confirmed by Aja Bončina, under the supervision of Elena Buzan from the University of Primorska. The specimen was hunted during a regular hunting activity in the hunting ground with special purposes (LPN) Pohorje (location Jurgovo), Slovenia. No sampling permits were required. The specimen’s tissues (muscle, all inner organs, brain, testis, skin) were snap-frozen immediately after harvesting and stored in liquid nitrogen until DNA extraction.

### Vouchering information

Physical reference materials for the sequenced specimen have been deposited in the Slovenian Museum of Natural History, mammal collection (zbirka sesalcev) under the accession number PMSL31444.

Frozen reference tissue material of muscle and kidney is available from the same individual at the Biobank Slovenian Museum of Natural History under the voucher IDs PMS_TIS_4.

### Data Availability

*R. rupicapra* and the related genomic study were assigned to Tree of Life identifier (ToLID) ‘mRupRup1’ and all sample, sequence, and assembly information are available under the umbrella BioProject ID PRJEB73452. The sample information is available at the following BioSample accessions: SAMEA112797446, SAMEA112797485, and SAMEA112797487. The genome assembly is accessible from the European Nucleotide Archive ENA integrated into the INSDC under accession number GCA_963981305.1 and the annotated genome is available through the Ensembl Rapid Release page (projects.ensembl.org/erga-bge). Sequencing data produced as part of this project are available from the ENA at the following accessions: ERX11550622, ERX13168315, ERX11550624, and ERX11550625. Documentation related to the genome assembly and curation can be found in the ERGA Assembly Report (EAR) document available at github.com/ERGA-consortium/EARs/tree/main/Assembly_Reports/Rupicapra_rupicapra/mRupRup1. Further details and data about the project are hosted on the ERGA portal at portal.erga-biodiversity.eu/organism/SAMEA112797440.

### Genetic Information

The estimated genome size is 3.2 Gb, while the estimation based on reads kmer profiling is 2.6 Gb. This is a diploid genome with a haploid number of 29 chromosomes (2n=58), including XY sex chromosomes in males. This information for this species was retrieved from Genomes on a Tree (Challis et al., 2023).

### DNA/RNA processing

DNA was extracted from spleen tissue using the Blood & Cell Culture DNA Midi Kit (Qiagen) following the manufacturer’s instructions. DNA quantification was performed with a Qubit dsDNA BR Assay Kit (Thermo Fisher Scientific), and DNA integrity was assessed using a Femtopulse system (Genomic DNA 165 Kb Kit, Agilent). DNA was stored at +4ºC until used.

RNA was extracted from brain tissue using an RNeasy Blood & Tissue Mini Kit (Qiagen) according to the manufacturer’s instructions. RNA quantification was performed with the Qubit RNA BR Kit, and RNA integrity was assessed using a Bioanalyzer system (Eukaryote Total RNA Nano Kit, Agilent). RNA was stored at -80ºC until used.

### Library Preparation and Sequencing

For long-read whole genome sequencing, a library was prepared using the SQK-LSK114 Kit (Oxford Nanopore Technologies, ONT), which was then sequenced on a PromethION 24 A Series instrument (ONT). A short-read whole genome sequencing library was prepared using the KAPA Hyper Prep Kit (Roche). A Hi-C library was prepared from kidney tissue using the Dovetail Omni-C Kit (Cantata Bio), followed by the KAPA Hyper Prep Kit for Illumina sequencing (Roche). The RNA library was prepared using the KAPA mRNA Hyper prep kit (Roche). The short-read libraries were sequenced on a NovaSeq 6000 instrument (Illumina). In total, 125x Oxford Nanopore, 84x Illumina WGS shotgun, and 90x HiC data were sequenced to generate the assembly.

### Genome Assembly Methods

The genome was assembled using the CNAG CLAWS pipeline (Gomez-Garrido, 2024). Briefly, reads were preprocessed for quality and length using Trim Galore v0.6.7 and Filtlong v0.2.1, and initial contigs were assembled with NextDenovo v2.5.0, followed by polishing of the assembled contigs using HyPo v1.0.3, removal of retained haplotigs with purge-dups v1.2.6 and scaffolding with YaHS v1.2a. Finally, assembled scaffolds were curated via manual inspection using Pretext v0.2.5 with the Rapid Curation Toolkit (gitlab.com/wtsi-grit/rapid-curation) to remove any false joins and incorporate any sequences not automatically scaffolded into their respective locations in the chromosomal pseudomolecules (or super-scaffolds). Finally, the mitochondrial genome was assembled as a single circular contig of 16,433 bp using the FOAM pipeline (github.com/cnag-aat/FOAM) and included in the released assembly (GCA_963981305.1). Summary analysis of the released assembly was performed using the ERGA-BGE Genome Report ASM Galaxy workflow (De Panis, 2024b), incorporating tools such as BUSCO v5.5, Merqury v1.3, and others (see reference for the full list of tools).

### Genome Annotation Methods

A gene set was generated using the Ensembl Gene Annotation system (Aken et al., 2016), primarily by aligning short-read RNA-seq data from BioSample SAMEA112797485 to the genome. Gaps in the annotation were filled via protein-to-genome alignments of a select set of vertebrate proteins from UniProt (The UniProt Consortium, 2019), which had experimental evidence at the protein or transcript level. At each locus, data were aggregated and consolidated, prioritising models derived from RNA-seq data, resulting in a final set of gene models and associated non-redundant transcript sets. To distinguish true isoforms from fragments, the likelihood of each open reading frame (ORF) was evaluated against known vertebrate proteins. Low-quality transcript models, such as those showing evidence of fragmented ORFs, were removed. In cases where RNA-seq data were fragmented or absent, homology data were prioritised, favouring longer transcripts with strong intron support from short-read data. The resulting gene models were classified into three categories: protein-coding, pseudogene, and long non-coding. Models with hits on known proteins and few structural abnormalities were classified as protein-coding. Models with hits to known proteins but displaying abnormalities, such as the absence of a start codon, non-canonical splicing, unusually small intron structures (<75 bp), or excessive repeat coverage, were reclassified as pseudogenes. Single-exon models with a corresponding multi-exon copy elsewhere in the genome were classified as processed (retrotransposed) pseudogenes. Models that did not fit any of the previously described categories, did not overlap protein-coding genes, and were constructed from transcriptomic data were considered potential lncRNAs. Potential lncRNAs were further filtered to remove single-exon loci due to their unreliability. Putative miRNAs were predicted by performing a BLAST search of miRBase (Kozomara et al., 2019) against the genome, followed by RNAfold analysis (Gruber et al., 2008). Other small non-coding loci were identified by scanning the genome with Rfam (Kalvari et al., 2018) and passing the results through Infernal (Nawrocki & Eddy, 2013). Summary analysis of the released annotation was carried out using the ERGA-BGE Genome Report ANNOT Galaxy workflow (De Panis, 2024a), incorporating tools such as BUSCO v5.5, OMArk 0.3 and others (see reference for the full list of tools).

## Results

### Genome Assembly

The genome assembly had a total length of 2,623,004,604 bp in 76 scaffolds including the mitogenome (Figures 1 and 2), with a GC content of 41.98%. It featured a contig N50 of 77,662,214 bp (L50=13) and a scaffold N50 of 100,920,001 bp (L50=10). There were 48 gaps, totaling 9,600 kb in cumulative size. The single-copy gene content analysis using the Mammalia database with BUSCO (Manni et al., 2021) resulted in 96.0% completeness (93.6% single and 2.4% duplicated). Additionally, 97.3% of reads k-mers were present in the assembly and the assembly has a base accuracy Quality Value (QV) of 55 as calculated by Merqury (Rhie et al., 2020).

**Figure 1.**
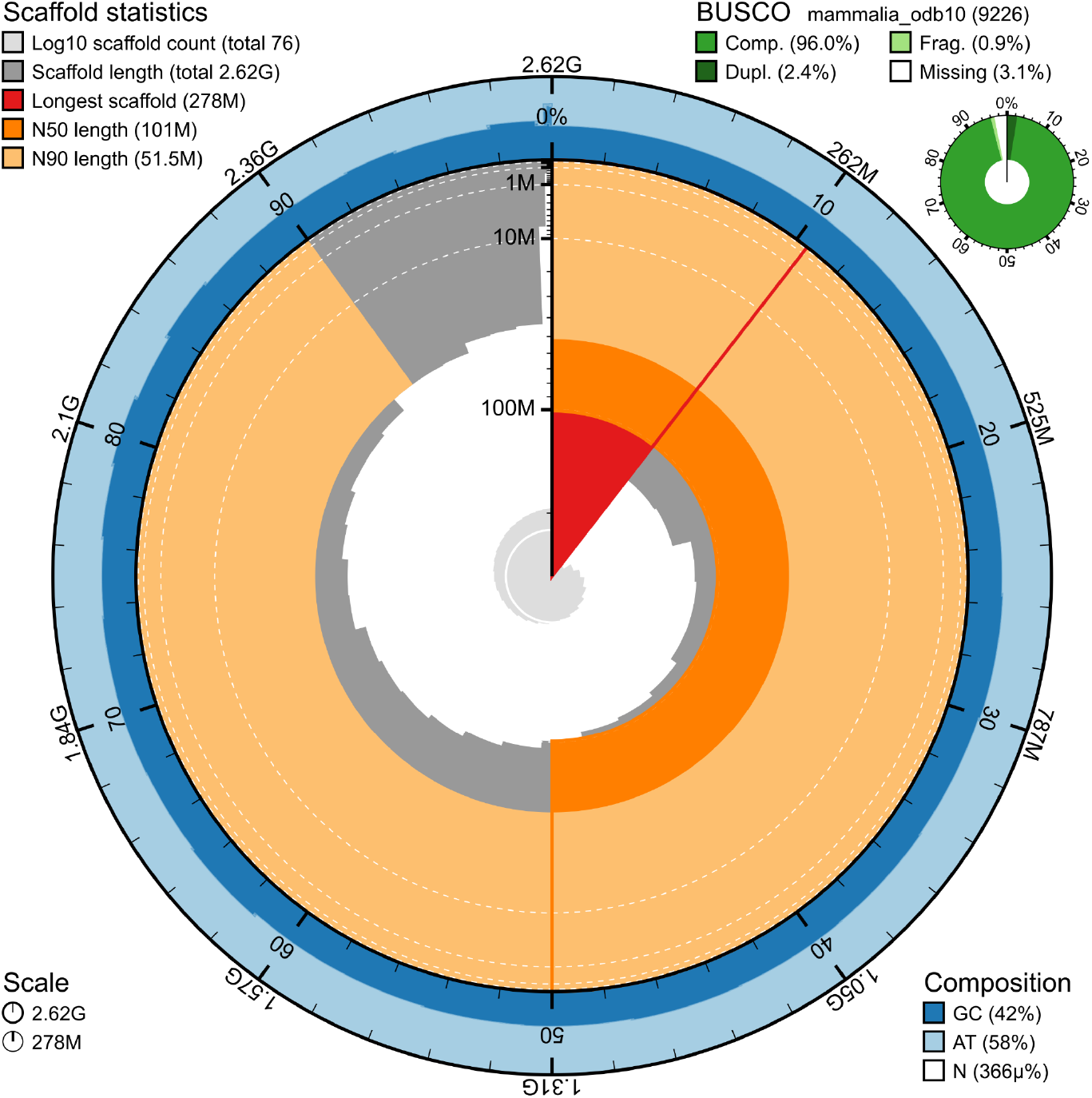
Snail plot summary of assembly statistics. The main plot is divided into 1,000 size-ordered bins around the circumference, with each bin representing 0.1% of the 2,623,004,604 bp assembly including the mitochondrial genome. The distribution of sequence lengths is shown in dark grey, with the plot radius scaled to the longest sequence present in the assembly (277,742,375 bp, shown in red). Orange and pale-orange arcs show the scaffold N50 and N90 sequence lengths (100,920,001 and 51,534,373 bp), respectively. The pale grey spiral shows the cumulative sequence count on a log-scale, with white scale lines showing successive orders of magnitude. The blue and pale-blue area around the outside of the plot shows the distribution of GC, AT, and N percentages in the same bins as the inner plot. A summary of complete, fragmented, duplicated, and missing BUSCO genes found in the assembled genome from the Mammalia database (odb10) is shown on the top right.

**Figure 2.**
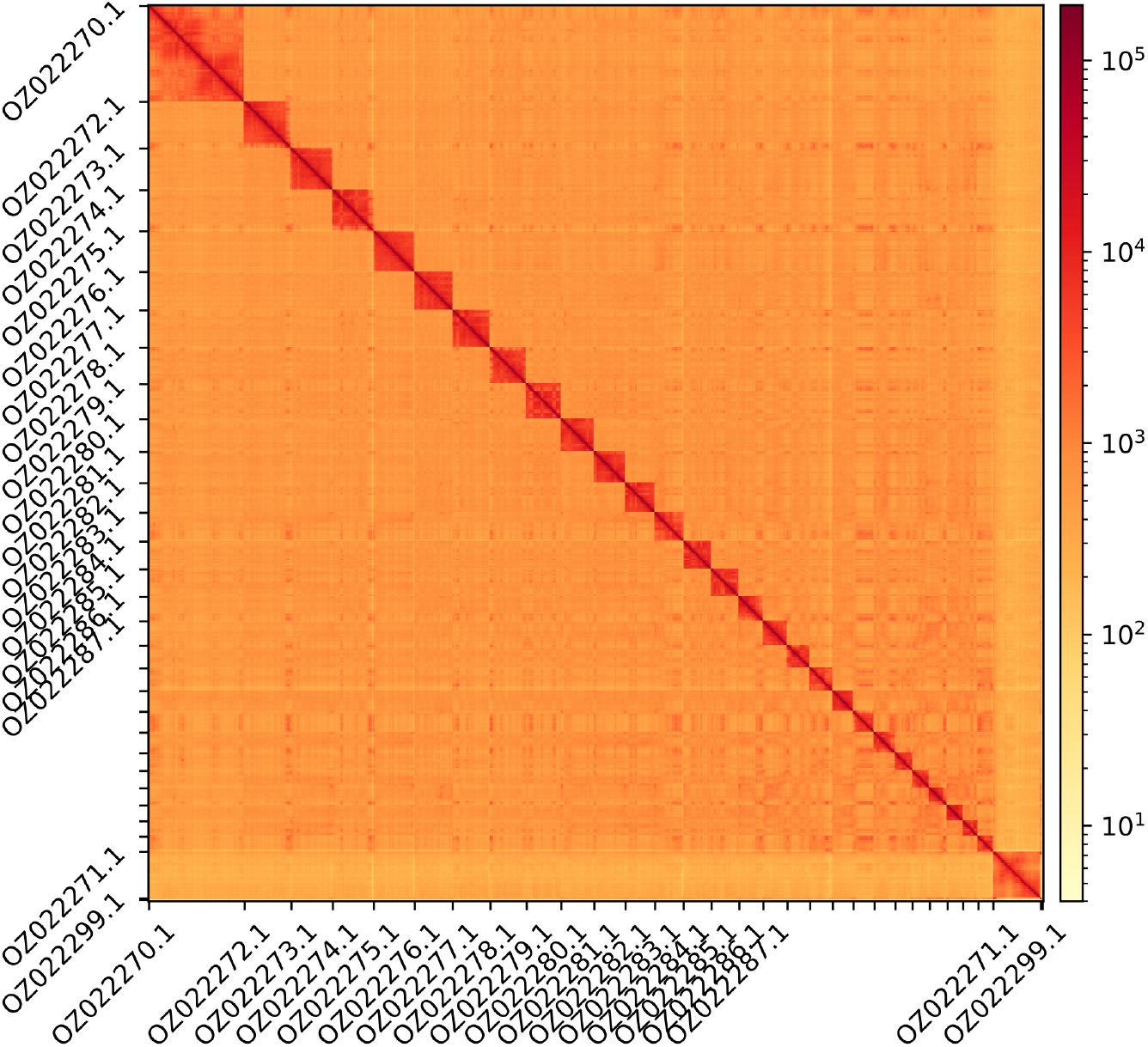
Hi-C contact map showing spatial interactions between regions of the genome. The diagonal corresponds to intra-chromosomal contacts, depicting chromosome boundaries. The frequency of contacts is shown on a logarithmic heatmap scale. Hi-C matrix bins were merged into a 50 kb bin size for plotting. Due to space constraints on the axes, only the GenBank names of the 17th largest autosomes and the sex chromosomes are shown.

### Genome Annotation

The genome annotation consisted of 18,339 protein-coding genes with an associated 30,055 transcripts, in addition to 3,032 non-coding genes (Table 1). Using the longest isoform per transcript, the single-copy gene content analysis using the Mammalia database with BUSCO resulted in 96.6% completeness. Using the OMAmer Artiodactyla database for OMArk (Nevers et al., 2024) resulted in 98.18% completeness and 98.83% consistency (Table 2).

**Table 1.** Statistics from assembled gene models.

|  | No. genes | No. transcripts | Mean gene length (bp) | No. single-exon genes | Mean exons per transcript |
| --- | --- | --- | --- | --- | --- |
| **mRNA** | 18,339 | 30,055 | 994,738,804 | 1,665 | 11.5 |
| **pseudogene** | 564 | 564 | 14,782 | 48 | 10.3 |
| **snoRNA** | 670 | 670 | 114 | 670 | 1.0 |
| **lncRNA** | 222 | 370 | 32,926 | 19 | 3.3 |
| **miRNA** | 215 | 215 | 89 | 215 | 1.0 |
| **snRNA** | 1,255 | 1,255 | 113 | 1,255 | 1.0 |
| **rRNA** | 59 | 59 | 189 | 59 | 1.0 |
| **scRNA** | 28 | 28 | 164 | 28 | 1.0 |
| **Other ncRNA** | 19 | 19 | 291 | 19 | 1.0 |

**Table 2.** Annotation completeness and consistency scores calculated by BUSCO run in protein mode (Mammalia) and OMArk (Artiodactyla)

|  | Complete | Singular | Duplicated | Fragmented | Missing |
| --- | --- | --- | --- | --- | --- |
| **BUSCO** | 8,915 (96.6%) | 8,828 (95.7%) | 87 (0.9%) | 39 (0.4%) | 272 (3.0%) |
| **OMArk** | 12,812 (98.18%) | 12,588 (96.46%) | 224 (1.72%) | - | 238 (1.82%) |
|  | Consistent | Inconsistent | Contaminants | Unknown |  |
| **OMArk** | 18,269 (98.83%) | 149 (0.81%) | 0 (0.00%) | 67 (0.36%) |  |

## Acknowledgements

We would like to express our gratitude to the Hunters Association of Slovenia and the Slovenia Forest Service (especially gamekeepers in LPN Pohorje), who helped in the sampling procedure as well as established a unique and important database supporting wildlife conservation and research in Slovenia. Special thanks are due to the Ministry of Agriculture, Forestry and Food for supporting wildlife research, including our sample collection. We acknowledge the assembly reviewer, Jonathan Wood, from the Wellcome Trust Sanger Institute.

## Conflict of Interest

The authors declare no conflict of interest related to this study. The funding sources had no involvement in the study design, collection, analysis, or interpretation of data; in the writing of the manuscript; or in the decision to submit the article for publication. All authors have participated sufficiently in the work to take public responsibility for the content and agree to the submission of this manuscript.

## Funder Information

Biodiversity Genomics Europe **(Grant no.101059492)** is funded by Horizon Europe under the Biodiversity, Circular Economy and Environment call (REA.B.3); co-funded by the Swiss State Secretariat for Education, Research and Innovation (SERI) under contract numbers 22.00173 and 24.00054; and by the UK Research and Innovation (UKRI) under the Department for Business, Energy and Industrial Strategy’s Horizon Europe Guarantee Scheme.

## Author Contributions

EB coordinated the project, BP collected the species, EB, AjB identified the species, EB, BP, and AjB sampled and preserved biological material and provided metadata, RM, NE, RF, and AsB provided support in sampling, shipping of biological material, metadata collection, and management, MG extracted DNA, prepared libraries, and performed sequencing, FCF, JGG and FC performed genome assembly and curation under the supervision of TSA, LH and FM performed genome annotation, DDP generated the analysis and report. All authors contributed to the writing, review, and editing of this genome note and read and approved the final version.

## Notes

### Competing Interest Statement

The authors have declared no competing interest.

